# Whole-mount three-dimensional imaging and quantitative analysis platform for early brain development using dorsal forebrain spheroids

**DOI:** 10.64898/2026.11.14.766664

**Authors:** Amara Ayoub, Lucy Lu, Tarik F Haydar

## Abstract

Three-dimensional culture systems have emerged as a bridge between *in vitro* and *in vivo* models. Organoids and spheroids derived from human stem cells can model cellular interactions, morphology and organization of the neocortex, and provide a platform for future translational research into neurological disorders. However, the heterogeneity commonly found in 3D culture systems presents considerable challenges for identifying significant, reproducible, and disorder-relevant phenotypes. Here, we present a rigorous methodological framework for the longitudinal analysis of dorsal forebrain spheroids, beginning with gross spheroid morphology, extending to structural phenotypes in rosette morphogenesis, and culminating in cellular analyses of neural progenitor cell proliferation and cell cycle dynamics. This workflow integrates isogenic iPSC-derived spheroid culture, whole-mount optical clearing, high-resolution 3D imaging and reconstruction, and cumulative EdU-labeling assays. Testing the sensitivity of this approach in an isogenic Down syndrome model, we reliably detected new features of altered spheroid growth, delayed rosette emergence, reduced rosette density, and prolonged progenitor cell cycle progression. This approach provides a reproducible strategy for measuring complex developmental phenotypes in 3D culture models and can be applied to neurodevelopmentally altered cell lines beyond Down syndrome.

## INTRODUCTION

Human cerebral organoids have emerged as an experimentally tractable *in vitro* representation of human cortical development (Birey et al., 2017; Kadoshima et al., 2013; Lancaster et al., 2013; Li et al., 2017). Organoids are complex 3D structures generated from stem or progenitor cells, including induced pluripotent stem cells (iPSCs) (Lancaster et al., 2013). These culture systems can recapitulate the cellular composition of ventricular and subventricular progenitor zones and ventricle-like cavities of the neocortex (Kadoshima et al., 2013; Lancaster et al., 2013; Li et al., 2017). Therefore, human cellular models offer a unique opportunity to study the developmental mechanisms underlying human-specific traits, such as diverse progenitor cell populations governing brain growth and neocortical expansion (Camp et al., 2015; Lui et al., 2011).

Organoids can contain areas representing multiple brain-region identities within a single complex structure, in which case they are commonly referred to as cerebral organoids (Di Lullo and Kriegstein, 2017; Lancaster et al., 2013). The heterogeneity in cerebral organoid systems presents considerable challenges for identifying significant, reproducible and disorder-relevant phenotypes (Heide et al., 2018; Sandoval et al., 2024). Specifically, organoids assembled from the same cell line under seemingly identical conditions can produce structures with different regional identities, adding substantial spatial and cellular heterogeneity, which can result in batch-to-batch variability between organoids (Heide et al., 2018; Sandoval et al., 2024). This is especially true for self-patterned organoids undergoing self-assembly with minimal addition of extrinsic factors (Heide et al., 2018). In contrast, region-specific spheroids are generated based on controlled spatiotemporal delivery of patterning factors that guide differentiation of cells toward a specific regional fate, such as forebrain spheroids (Birey et al., 2017; Yoon et al., 2019). Here we provide a workflow that employs patterned spheroids to comprehensively measure cortical development with high reproducibility and specificity. In particular, we show how these methods can uncover novel understanding of the cellular and molecular processes of brain growth in Down syndrome (DS).

Cortical spheroids commence with a pseudo-neurulation event generating neural rosettes that resemble the neural tube primordium *in vivo* (Miotto et al., 2023; Zhang et al., 2001). Neural rosettes form when neural progenitor cells (NPCs) polarize and express adhesion molecules, forming an apical surface. This apical band then becomes circularized and tethered by adhesion molecules, forming a central lumen. Neural rosettes act as proliferative morphogenetic centers of the spheroid, guiding spheroid development and growth (Harding et al., 2014; Knight et al., 2018; Miotto et al., 2023). Current characterization methods often sacrifice spatial information through the use of two-dimensional (2D) histological analyses, leaving the morphogenetic details enabled by whole-tissue investigation underexplored (Albanese et al., 2020). As rosettes vary in size, shape, organization and position within the spheroid, analyses of selected 2D histological sections may underestimate rosette abundance, obscure their spatial context, exclude structures outside the sectioning plane, and preferentially sample structures with well-defined morphology. Therefore, methods that preserve the intact spheroid are needed to characterize the architectonic features, size, and cellular composition of rosettes throughout the entire spheroid. Comprehensive imaging and quantification of all rosettes provides a more detailed and representative analysis of their abundance, morphology, and spatial organization while capturing the full range of rosette structures present within the tissue. However, imaging intact spheroids remains technically challenging due to the reduced antibody and light penetration at greater tissue depths, limiting the resolution of internal structures. Whole-mount imaging also requires the spheroid to remain stable during image acquisition. To address these challenges, we optimized staining and optical clearing of intact spheroids with a mounting approach that preserved spheroid morphology and position during high-resolution imaging.

In this work, we describe methods designed to address the quantitative variation found in spheroid cultures using dorsal forebrain spheroids (DFSs). The approach, which combines region-specific spheroid patterning with longitudinal analysis, whole-mount optical clearing, high-resolution 3D imaging, and cell cycle analysis, enables specific and repeatable measures of spheroid growth sufficient to distinguish alterations in key developmental processes. A major strength of the study is the use of an isogenic iPSC system, which enables direct testing of our methods’ sensitivity and allows phenotypic differences to be directly attributed to the genetic condition being modeled rather than confounding genetic background variation (Hibaoui et al., 2014; Maclean et al., 2012; Murray et al., 2015; Weick et al., 2013). Although we developed this framework using an isogenic trisomy 21 line, the methodological strategy is not restricted to this condition. Instead, it is intended to provide a common foundation for investigating early events in neurodevelopmentally altered cell lines where changes in spheroid growth, tissue organization, NPC dynamics, or transcriptional programs may be subtle, heterogeneous, or developmentally dynamic (Czerminski et al., 2022; Gough et al., 2020; Sandoval et al., 2024).

### Rationale for a Rigorous 3D Culture Methodology

A major focus of this work is the development of a robust and rigorous experimental design that can quantitatively resolve early developmental phenotypes despite the line-to-line and batch-to-batch variability of organoid and spheroid systems. To address intrinsic variability, we used large sample sizes and longitudinal analysis to enable robust detection of phenotypic differences.

The framework was organized around three levels of analysis. First, longitudinal phase-contrast imaging was used to quantify spheroid growth over the first month in culture. Second, whole-mount immunostaining, optical clearing, multiphoton imaging and 3D reconstruction were used to quantify neural rosette emergence, organization, density and morphometry across intact spheroids over time. Third, PH3 quantification and cumulative EdU labeling were used to assess proliferative activity and cell-cycle progression within rosette NPCs. Together, this design enabled analysis across gross, structural and cellular levels of development.

**Table 1.** Key design elements.

| **Design element** | **Problem addressed** | **Implementation in this study** |
|---|---|---|
| Isogenic iPSC comparison | Confounding genetic background variation | Euploid and trisomic lines genetically identical apart from HSA21 triplication |
| Independent differentiations | Batch-to-batch variability | Three differentiations per genotype analyzed |
| Large sample sizes | Subtle, heterogeneous phenotypes | 24-48 spheroids per genotype and timepoint for growth analysis. Approximately 20-150 rosettes per single spheroid, depending on timepoint |
| Longitudinal imaging | Cross-sectional analyses do not address developmental timing or changes within spheroids over time | Spheroids analyzed consecutively over the first month in culture |
| Whole-mount clearing and 3D imaging | Loss of spatial information in sectioned tissue | Intact spheroids stained, cleared, imaged and reconstructed in 3D |
| Distribution-based analysis | Mean-based analysis can obscure subpopulations | Gaussian mixture modeling and stratification were used to assess population-level heterogeneity |
| Rosette scoring | Inconsistent inclusion of abnormal structures | Rosettes classified based on lumen continuity, radial alignment, polarity and morphology |
| Cumulative EdU labeling | Transient proliferation markers alone are incomplete | EdU labeling used to estimate growth fraction, total cell cycle length, S-phase duration and the other components of the cell cycle |

## RESULTS

Our general approach was to apply our analytical methodology to characterize developmental trajectories and heterogeneity within control, diploid DFSs. We then used an isogenic trisomy 21 line to determine whether these methods could resolve genotype-associated differences from the euploid developmental trajectory. This workflow allowed us to define features of spheroid development and evaluate its ability to distinguish altered developmental patterns associated with trisomy 21.

### Longitudinal analysis defines spheroid growth trajectories and heterogeneity

DFSs were generated in accordance with established protocols (Birey et al., 2017; Yoon et al., 2019; **Figures 1A-D**). To characterize DFS growth, a longitudinal time-course analysis of spheroid growth and size was performed using phase-contrast imaging. Individual spheroids were tracked every 2 days from 8 to 30 days in vitro (DIV), with 24-48 spheroids analyzed at each time point across three independent differentiations. Phase-contrast images of live whole spheroids were captured using a Keyence BZ-X710 inverted fluorescence microscope (**Figures 1E, F**). Spheroid area was measured using QuPath image analysis software following pixel classification-based segmentation, whereby annotations were generated to delineate spheroid boundaries and calculate projected two-dimensional area. Given the variability in spheroid size both across differentiations and between genotypes, large sample sizes and longitudinal analysis were implemented to accurately resolve subtle but biologically relevant differences in growth dynamics. An a priori power analysis using variance estimates from our pilot study indicated that 20 samples per genotype and timepoint were required to detect the observed effect size with 80% power at an alpha level of 0.05.

**Figure 1.**
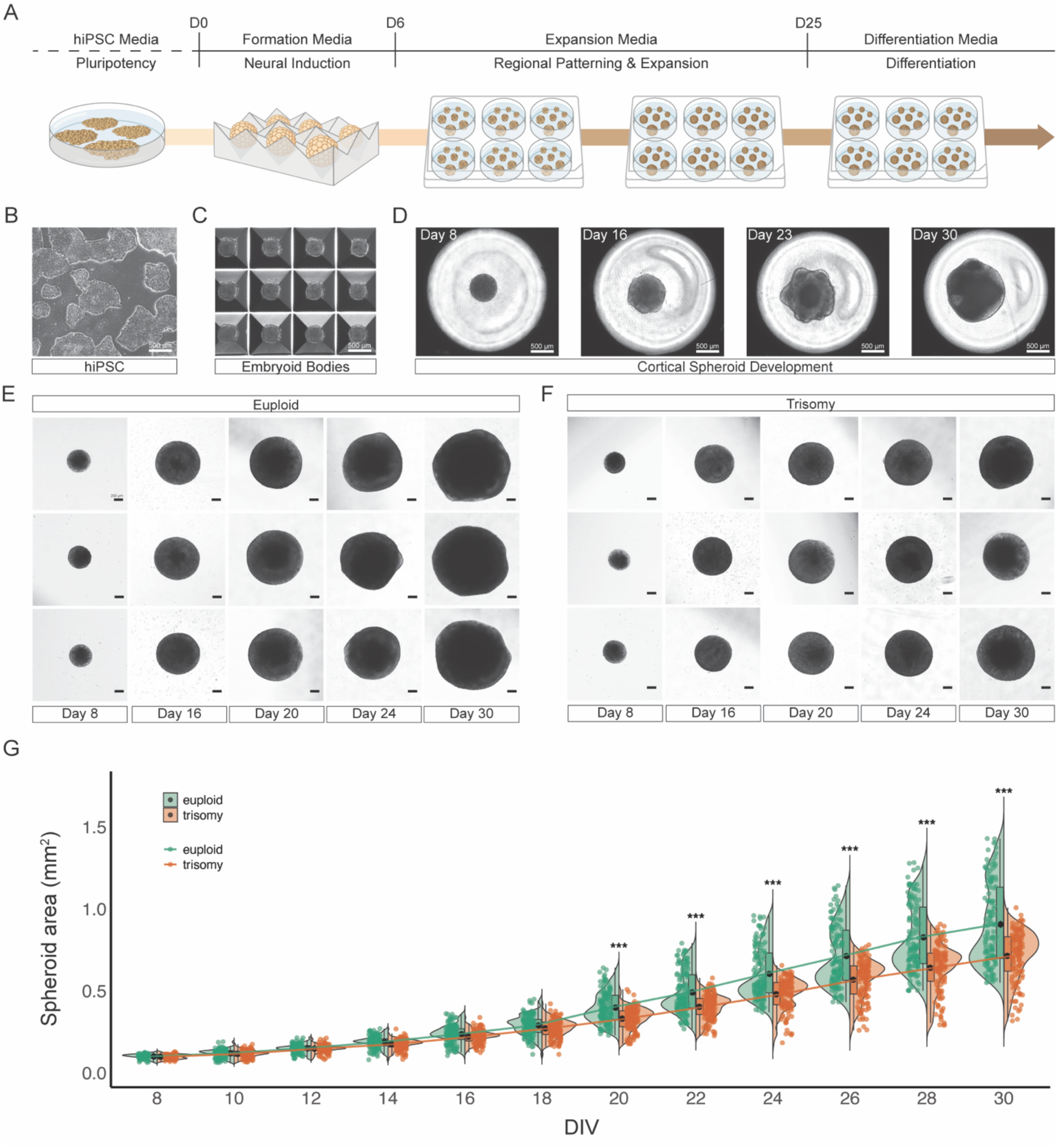
Generation and growth characterization of dorsal forebrain spheroids. **(A)** Schematic of the dorsal forebrain spheroid (DFS) culture system. **(B-D)** Representative phase contrast images at each stage of differentiation. **(B)** iPSC colonies maintained in two-dimensional culture prior to aggregation. **(C)** Embryoid bodies generated within a microwell culture system. **(D)** DFS morphogenesis at 8, 16, 23, and 30 DIV, showing progressive increase in spheroid size and structural complexity over time. Scale bars, 500 μm. **(E-F)** Representative phase contrast images of individual euploid **(E)** and trisomic **(F)** DFSs across early developmental timepoints. Scale bars, 200 μm. **(G)** Longitudinal quantification of spheroid area from 8 to 30 DIV in euploid and trisomic DFSs. Individual data points represent individual spheroids, violin plots show the distribution of spheroid areas at each timepoint, and overlaid lines represent genotype-specific growth trajectories. Box-plots indicate the 25^th^ to 75^th^ percentiles, with the mean shown as a black dot. Euploid and trisomic spheroids exhibited comparable growth from 8-18 DIV, followed by significant divergence beginning at 20 DIV, with trisomic spheroids displaying reduced area relative to euploid controls. DIV: days in vitro. Data collected from three independent differentiations, with 24-48 spheroids per genotype per timepoint. Data presented as mean ± standard error (SE). Statistical significance indicated as \*\*\**p* ≤ 0.001.

Spheroids exhibited progressive growth over time, consistent with the expected expansion dynamics reported for cortical spheroids (**Figure 1E, G**). Repeated measurements across development enabled characterization of growth trajectories both within and across independent differentiations. Although the population of spheroids showed a consistent increase in area over time, individual spheroid measurements revealed group variability at each timepoint. To examine the distribution underlying this variability, size distribution, mean spheroid area, and individual spheroid measurements were assessed at each developmental timepoint. Rather than forming a single unimodal Gaussian distribution, spheroid measurements exhibited multiple distributional peaks across each of the three independent differentiations from 20-30 DIV (**Figure 2A-D**). Gaussian mixture modeling identified predominantly bimodal distributions consistent with distinct spheroid size subpopulations within each genotype (**Figure 2E**). The lower- and upper-mean Gaussian components were designated as small and large populations, respectively. Euploid spheroids remained significantly larger than trisomic spheroids within both groups from 20-30 DIV (**Figure 2F**). These distributional differences would not be captured by analyses based solely on average growth metrics, highlighting the importance of large sample sizes for resolving population-level heterogeneity in 3D culture systems.

**Figure 2.**
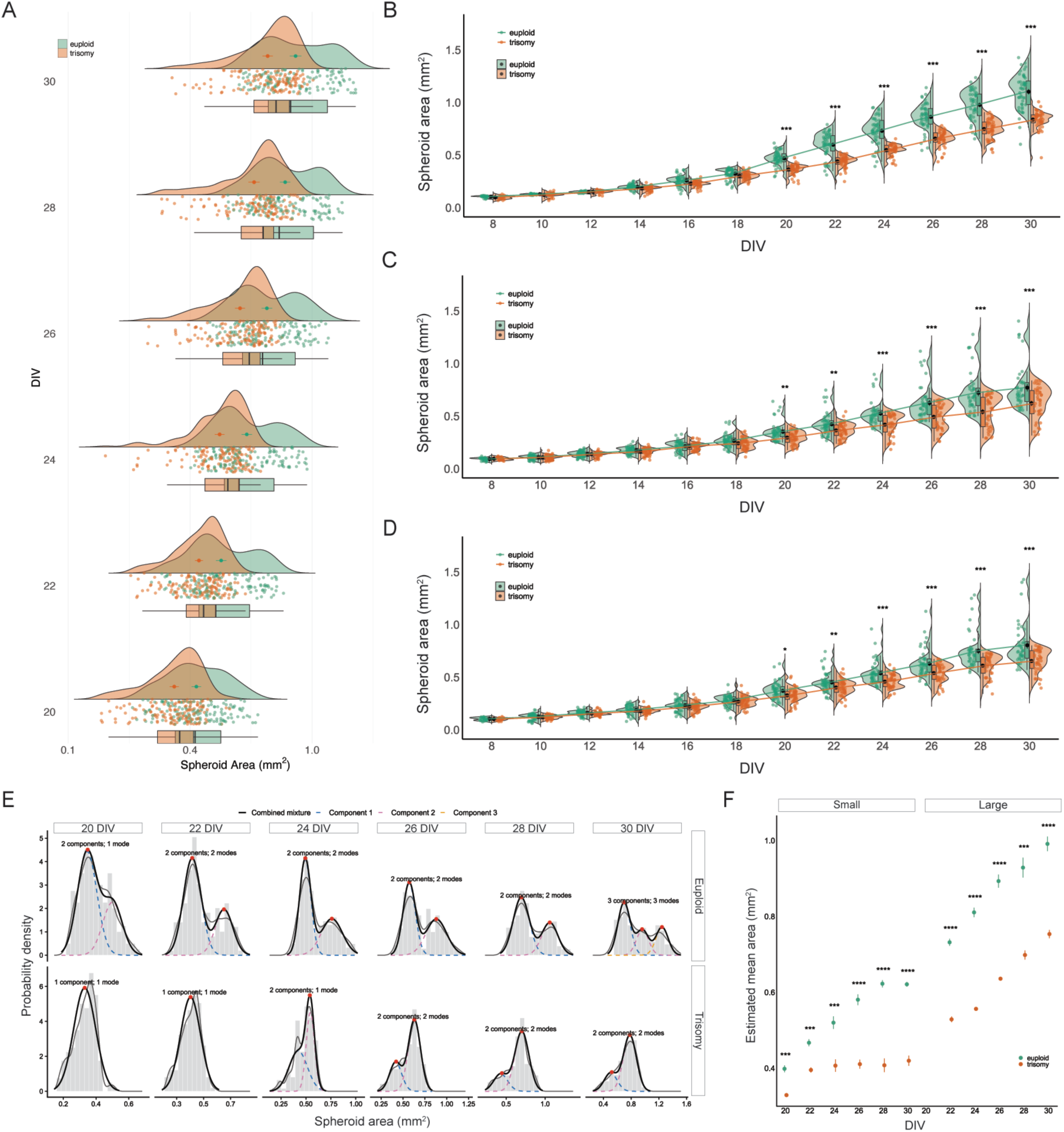
Size distribution analyses of DFS subpopulations. (**A**) Raincloud plot of spheroid area distributions from 20 to 30 DIV for pooled euploid and trisomic DFSs across all three differentiations. Individual data points represent individual spheroids, half-violin plots show the distribution of spheroid areas at each timepoint, and boxplots indicate interquartile range. Both genotypes exhibit heterogeneous and multimodal size distributions over time. **(B-D)** Longitudinal quantification of spheroid area from 8 to 30 DIV for differentiation one **(B)**, two **(C)**, and three **(D)**. Individual data points represent individual spheroids, violin plots show the distribution of spheroid areas at each timepoint, and overlaid lines represent genotype-specific growth trajectories. In each independent differentiation, euploid spheroids exhibited a larger average area than trisomic spheroids over time. Data presented as mean ± SE. **(E)** Gaussian mixture modeling of spheroid area distributions from 20 to 30 DIV, performed separately by genotype and timepoint. Dashed curves represent the individual fitted components, solid black curves show the overall fitted distributions, and red points mark the fitted modes. The number of components was selected using the Bayesian information criterion. **(F)** Comparison of small and large subpopulations, defined by the lower- and upper-mean Gaussian components, respectively. Euploid spheroids remained significantly larger than trisomic spheroids within both populations. Data presented as estimated mean ± SE. Statistical significance indicated as \**p* ≤ 0.05, \*\**p* ≤ 0.01, \*\*\**p* ≤ 0.001, and \*\*\*\**p* ≤ 0.0001.

The same longitudinal and distribution-based analyses were then applied to trisomic DFSs. Euploid and trisomic DFSs developed comparably during early stages of development from 8-18 DIV, with no significant differences in spheroid area detected at any timepoint within this interval. Trisomic spheroids diverged from control in both growth rate and size beginning at 20 DIV and this growth divergence persisted and became more pronounced through 30 DIV (**Figure 1F, G**). The robust sample sizes enabled us to identify a significant shift in the distribution of spheroid sizes in trisomic cultures. Specifically, the spheroid size distributions revealed an increased representation of smaller trisomic spheroids at later developmental timepoints. Distribution-based analyses indicated that the overall reduction in mean trisomic spheroid area was driven by the persistence of a smaller subpopulation with limited expansion over time. In particular, these findings demonstrate that a population-level analysis can elucidate specific cellular features that may only be present in a subset of the biological sample.

This multimodal structure demonstrates that mean spheroid area alone does not fully describe the growth behavior of the underlying population. Together, these findings demonstrate that large sample sizes, longitudinal analysis, and distribution-based approaches can identify when growth phenotypes emerge and determine whether differences in mean spheroid size reflect a uniform population-wide change or the presence of distinct spheroid subpopulations.

### Whole-mount 3D imaging preserves rosette spatial organization

Neural rosettes are composed of radially organized Sox2^+^ NPCs that polarize around an Ncad^+^ central lumen, resembling the organization of the developing neural tube and its ventricular zone. As the proliferative units within spheroids, neural rosettes are the main focus of subsequent analyses in this work. Unlike previous studies of cryosectioned organoids, rosettes in this work were characterized in intact spheroids using a novel whole-mount optical clearing approach, preserving their spatial organization and enabling a wholistic set of cellular and spatial measurements. Whole-intact spheroids were immunostained for SRY-Box 2 (Sox2) and N-cadherin (Ncad) and imaged by multiphoton microscopy using stitched tile scans and high-resolution z-stacks. Image sets were then reconstructed in 3D to visualize rosette architecture throughout the entire spheroid volume (**Figure 3A**).

**Figure 3.**
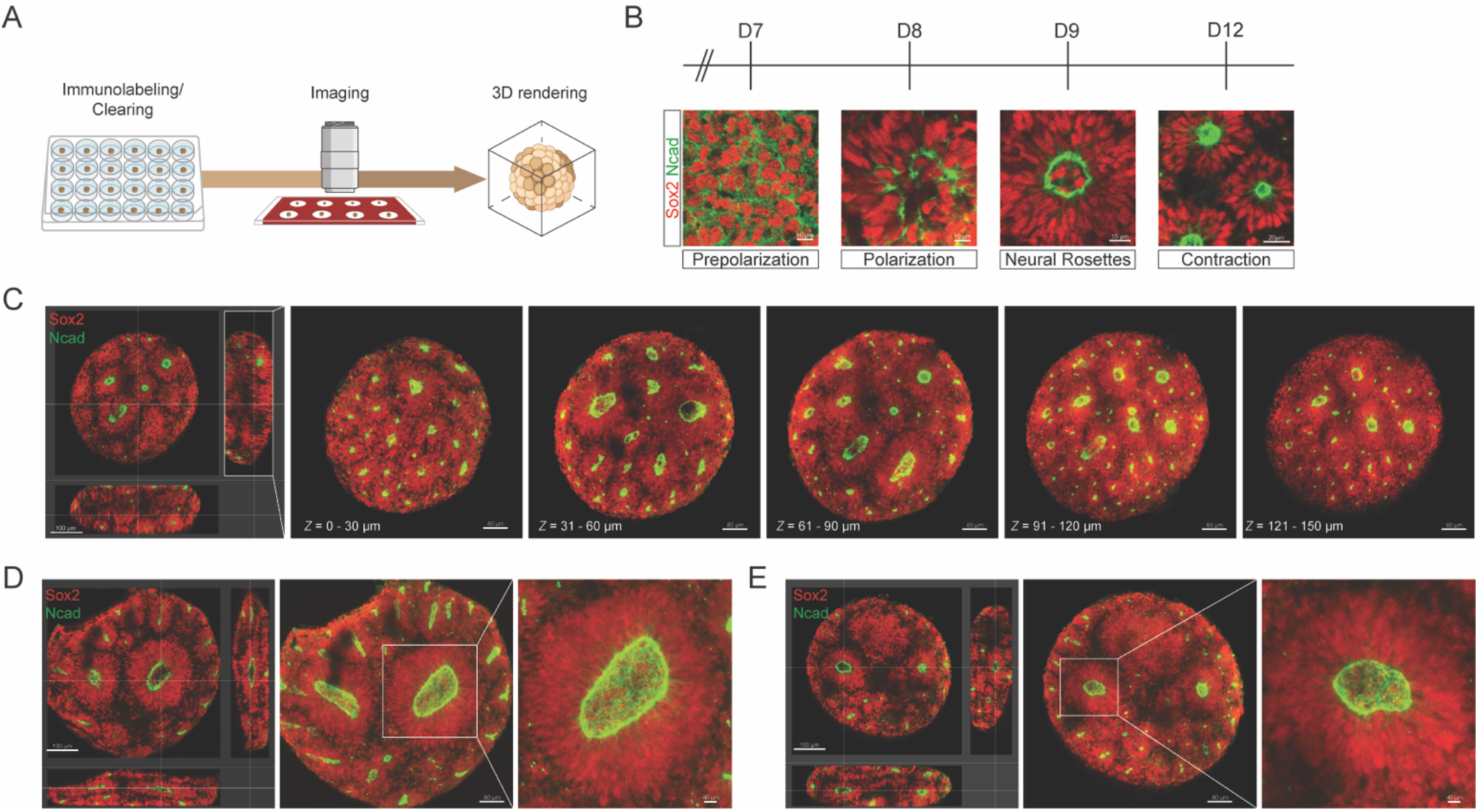
High-resolution 3D imaging reveals rosette heterogeneity. **(A)** Schematic of the staining, imaging and 3D reconstruction workflow used to visualize neural rosettes in dorsal forebrain spheroids. **(B)** Representative images of neural rosette morphogenesis from pre-polarization to contraction. Images illustrate sequential stages of rosette emergence, including early scattered adhesion molecule expression, polarization foci, neural rosette formation, and subsequent contraction of peripheral rosettes. **(C)** Orthogonal view and serial optical sections through a whole cleared spheroid shown at 30 μm intervals across a total depth of 150 μm. Each image represents a maximum-intensity projection of the indicated z-interval, demonstrating the distribution of rosettes within the spheroid. **(D-E)** Representative images of deep internal rosette structures in euploid **(D)** and trisomic **(E)** spheroids at 16 DIV. This illustrates distinct peripheral and internal rosette morphologies. Scale bars, as indicated. Sox2, red; Ncad, green.

Because the timing at which rosettes emerge can vary substantially across protocols and different cell lines, preliminary studies were conducted using our culture protocol to define the timing of neural rosettes emergence in DFSs. Consistent with previous reports (Knight et al., 2018), rosettes emerged through four distinct stages: pre-polarization, polarization foci, neural rosette, and contraction (**Figure 3B**). At 7 DIV, scattered expression of adherens molecules was evident amongst the Sox2^+^ progenitor cells. These cellular connections then formed polarization foci by 9 DIV as the progenitor cells began to align and consolidate around Ncad^+^ adhesion molecules. The neural rosette stage was marked by the formation of circular neuroepithelia, each containing a clear central lumen with strong adherens expression. By 12 DIV, neural rosettes on the outer surface of the spheroid contracted, exhibiting tighter Ncad^+^ lumens. Based on these observations, the onset of rosette formation was identified at 9 DIV. These observations established a reproducible sequence of rosette morphogenesis that could be used to comprehensively identify differences in the timing or organization of rosette development.

High-resolution 3D images of entire cleared spheroids were acquired at 1 μm Z-step intervals. A representative serial z-stack spanning a depth of 150 μm is shown in **Figure 3C**, with optical sections displayed at 30 μm spacing. This approach elucidated that rosette structures are present throughout the depth of the spheroid rather than being restricted to the surface or a single plane. Reconstruction of intact spheroids revealed heterogeneity in rosette morphology as well, with rosettes acquiring different shapes depending on their position. Rosettes located near the spheroid periphery were generally more spherical and contracted, with tighter Ncad^+^ lumens and compact Sox2^+^ NPC organization. In contrast, internal rosettes were larger and contained broader luminal structures surrounded by more expansive Sox2^+^ NPC regions (**Figures 3D, E**). Three-dimensional reconstruction preserved the position of these structures within the intact spheroid and demonstrated that rosette morphology varied across peripheral and internal spatial domains. Because internal rosettes may not be represented within a selected tissue section, their abundance and morphology would not be readily identified using traditional histological sectioning of spheroids.

### Rosette structural classification

Given the heterogeneity in rosette morphology, a structured scoring system was developed to establish consistent criteria for evaluation of rosette organization. Rosettes were scored based on Ncad+ lumen continuity, Sox2+ radial alignment, number of lumens, apical-basal polarity and overall morphology (**Table 2**). Using these criteria, rosettes were categorized as organized, disorganized, or disrupted. This classification scheme enabled consistent evaluation of rosettes in each organizational class across spheroids, developmental timepoints, and genotypes. It also distinguished changes in rosette organization from changes in the structural dimensions of organized rosettes. Only organized rosettes with clearly defined luminal and neuroepithelial boundaries were included in subsequent surface-based 3D morphometric analyses.

**Table 2.**
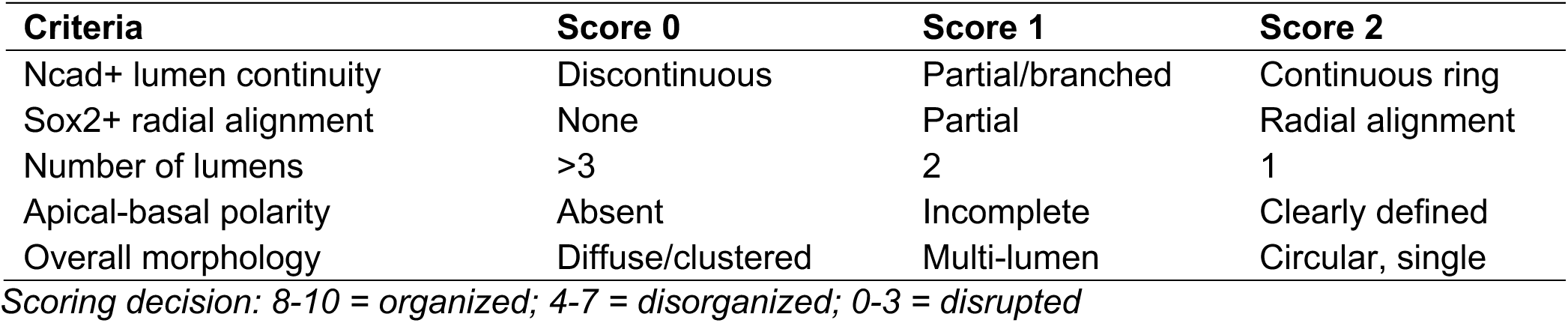
Rosette organization criteria and scoring system.

Representative example images of these organizational states are shown in **Figure 4A**. Organized rosettes exhibited a continuous Ncad^+^ lumen and radially aligned Sox2^+^ NPCs, whereas disorganized rosettes showed partial or branched lumen continuity together with altered radial organization. Disrupted rosettes lacked coherent lumen structure and exhibited diffuse or clustered Sox2^+^ progenitor organization. Absolute quantification of rosette categories confirmed that trisomic spheroids exhibited substantial organizational defects at 23 and 30 DIV (**Figure 4B**). These findings indicate that trisomic spheroids develop marked defects in rosette maintenance and structural integrity by 23 DIV, with disrupted rosettes representing the predominant abnormal category.

**Figure 4.**
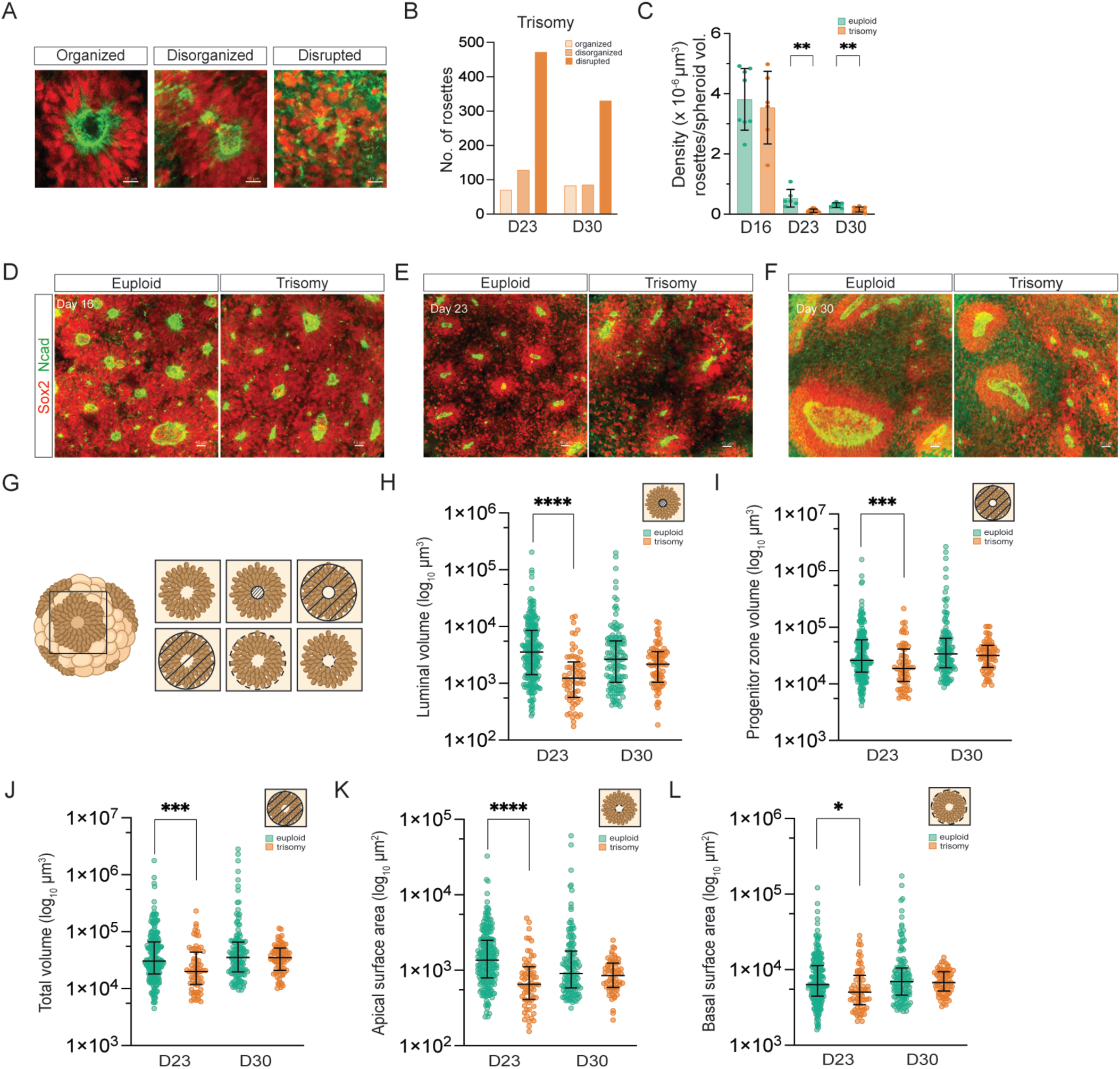
Rosette organization and density in DFSs. **(A)** Representative images of rosettes classified as organized, disorganized, and disrupted. (**B**) Absolute quantification of rosette organization categories in trisomic DFSs at 23 and 30 DIV. Trisomic spheroids exhibited a marked prevalence of disorganized and disrupted rosettes. (**C**) Quantification of rosette density (number of rosettes per spheroid volume) across developmental time points. Rosette density was comparable between genotypes at 16 DIV but significantly reduced in trisomic spheroids at 23 and 30 DIV. Density shown as mean ± standard deviation; unpaired Student’s *t*-test. (**D-F**) Representative images of euploid and trisomic spheroids at 16, 23, and 30 DIV. (**G**) Schematic illustrating the structural parameters used for 3D morphometric analysis of organized neural rosettes. (**H-L**) 3D morphometric quantification of individually segmented organized rosettes. Parameters include luminal volume (**H**), progenitor zone volume (**I**), total rosette volume (**J**), apical surface area (**K**), and basal surface area (**L**). Data collected from three independent differentiations, with 9 spheroids per genotype per timepoint. Each point represents a single organized rosette. Distributions shown as median and interquartile range; Type III ANOVA test. Statistical significance indicated as \**p* ≤ 0.05, \*\**p* ≤ 0.01, \*\*\**p* ≤ 0.001, and \*\*\*\**p* ≤ 0.0001. For parameters requiring well-defined boundaries, only organized rosettes were analyzed. Scale bars, as indicated. Sox2, red; Ncad, green.

Organized rosettes were quantified and normalized to the total spheroid volume to determine rosette density (**Figure 4C**). This approach enabled comparison of rosette abundance across development and genotypes while accounting for differences in spheroid size, revealing decreased rosette density in trisomic DFSs. Representative images of euploid and trisomic rosettes are shown in **Figure 4D-F**.

### 3D morphometry quantifies rosette size and architecture

To quantitatively examine the architectonic features, size and spatial organization of neural rosettes during spheroid development, high-resolution 3D images of intact cleared spheroids were reconstructed and segmented using Imaris analysis software. We first trained a machine learning segmentation model using three spheroids from each genotype to generate a reference dataset of surfaces, which identified and measured the majority of rosettes across the entire spheroid. Rosettes that were not accurately captured by this segmentation were then re-segmented individually within a user-defined region of interest. This semi-automated approach combines machine learning-based segmentation with manual refinement to accurately identify and quantify neural rosettes.

Rosette measurements were derived from nine spheroids per genotype at each timepoint, pooled across three spheroid replicates from three independent differentiations. A schematic of the structural parameters measured is shown in **Figure 4G**, including luminal volume, progenitor zone volume, total rosette volume, apical surface area, and basal surface area. Only luminal volume was quantified at 16 DIV, as rosettes overlapped substantially and lacked sufficiently discrete NPC boundaries for reliable segmentation of the full rosette architecture. Considerable heterogeneity was observed across rosette morphometric parameters (**Figures 4H-L**), with all measurements displaying right-skewed distributions within each genotype and timepoints.

Three-dimensional morphometric quantification of organized rosettes is shown in **Figures 4H-L**. Morphometric analyses first established how organized euploid rosettes changed between 23 and 30 DIV. Progenitor zone volume, total rosette volume, and basal surface area increased over time, consistent with the expansion of the Sox2^+^ NPC region. In contrast, luminal volume and apical surface area decreased, consistent with the contraction of the Ncad^+^ lumen during rosette development. Thus, control rosette maturation was characterized by expansion of the NPC compartment accompanied by contraction of the central lumen.

This morphometric analysis was then applied to characterize trisomic rosettes. Trisomic rosettes did not exhibit the same pattern found in euploid spheroids. In particular, both luminal volume and apical surface area were greater at 30 DIV than at 23 DIV. At 23 DIV, trisomic rosette values were reduced across all measured structural parameters relative to their euploid counterparts, including luminal volume (1,236 μm3, IQR = 1,813 μm3, trisomic vs 3,557 μm3, IQR = 7,158 μm3, euploid; *p* < 0.0001, **Figure 4H**), progenitor zone volume (18,627 μm3, IQR = 30,457 μm3, trisomic vs 25,935 μm3, IQR = 43,874 μm3, euploid; *p* < 0.001, **Figure 4I**), total rosette volume (19,904 μm3, IQR = 31,932 μm3, trisomic vs 30,466 μm3, IQR = 48,207 μm3, euploid; *p* < 0.001, **Figure 4J**), apical surface area (654 μm2, IQR = 704 μm2, trisomic vs 1,364 μm2, IQR = 1,713 μm2, euploid; *p* < 0.0001, **Figure 4K**) and basal surface area (5,036 μm2, IQR = 4,975 μm2, trisomic vs 6,346 μm2, IQR = 6,816 μm2, euploid; *p* < 0.05, **Figure 4L**). However, by 30 DIV trisomic rosettes were statistically comparable to euploid controls. These data showing that structural differences at 23 DIV became less pronounced as development proceeds, indicating that early structural defects may be compensated over time. Overall, these findings demonstrate that our 3D morphometric analyses carefully measure the structural maturation of euploid rosettes and are sufficient to elucidate the developmental alterations in trisomic rosette architecture that was most apparent at 23 DIV. We note that these structural abnormalities are coincident with the deflection in spheroid growth dynamics and that the normalization measured at 30 DIV does not correct the spheroid growth curve during this period.

### Measuring neural progenitor cell proliferation in rosettes

Rosette structures have been reported to replicate the cellular features of the pseudostratified neuroepithelium found in the embryonic and fetal nervous system (Miotto et al., 2023). To confirm this and to measure the basal proliferation rate in the rosettes, we first quantified the number of mitotic cells per rosette using phospho-histone H3 (PH3) staining. Whole-mount staining for PH3 and Ncad demonstrated that NPCs within organized rosettes were actively proliferative, with mitotic cells distributed around Ncad^+^ lumens, consistent with the expected apical localization of mitotic NPCs within the neuroepithelium (**Figure 5A**). We hypothesized that rosettes with greater luminal volume contain more proliferative NPCs. Therefore, we examined the relationship between mitotic activity and rosette size. PH3^+^ spots were computationally assigned to Ncad^+^ luminal surfaces using the spot-surface correlation function in Imaris software (**Figure 5A, inset**). In euploid DFSs, the number of PH3^+^ cells per rosette increased significantly over time (**Figure 5B**). In contrast, trisomic rosettes did not exhibit a significant increase in PH3^+^ cells per rosette over time (**Figure 5C**). Comparisons between genotypes revealed fewer PH3^+^ cells per rosette in trisomic spheroids across all timepoints, with differences most pronounced at 30 DIV (*p* <0.0001) (**Figure 5D**). Representative images of euploid and trisomic rosettes are shown in **Figure 5E-G**. Additionally, the number of PH3^+^ cells per rosette was positively associated with luminal volume in euploid DFSs, indicating that larger rosettes contained more mitotic NPCs. This relationship was weaker across all timepoints in trisomic DFSs (**Figure 5H-J**). This indicates that trisomy 21 attenuates NPC mitotic activity in rosettes.

**Figure 5.**
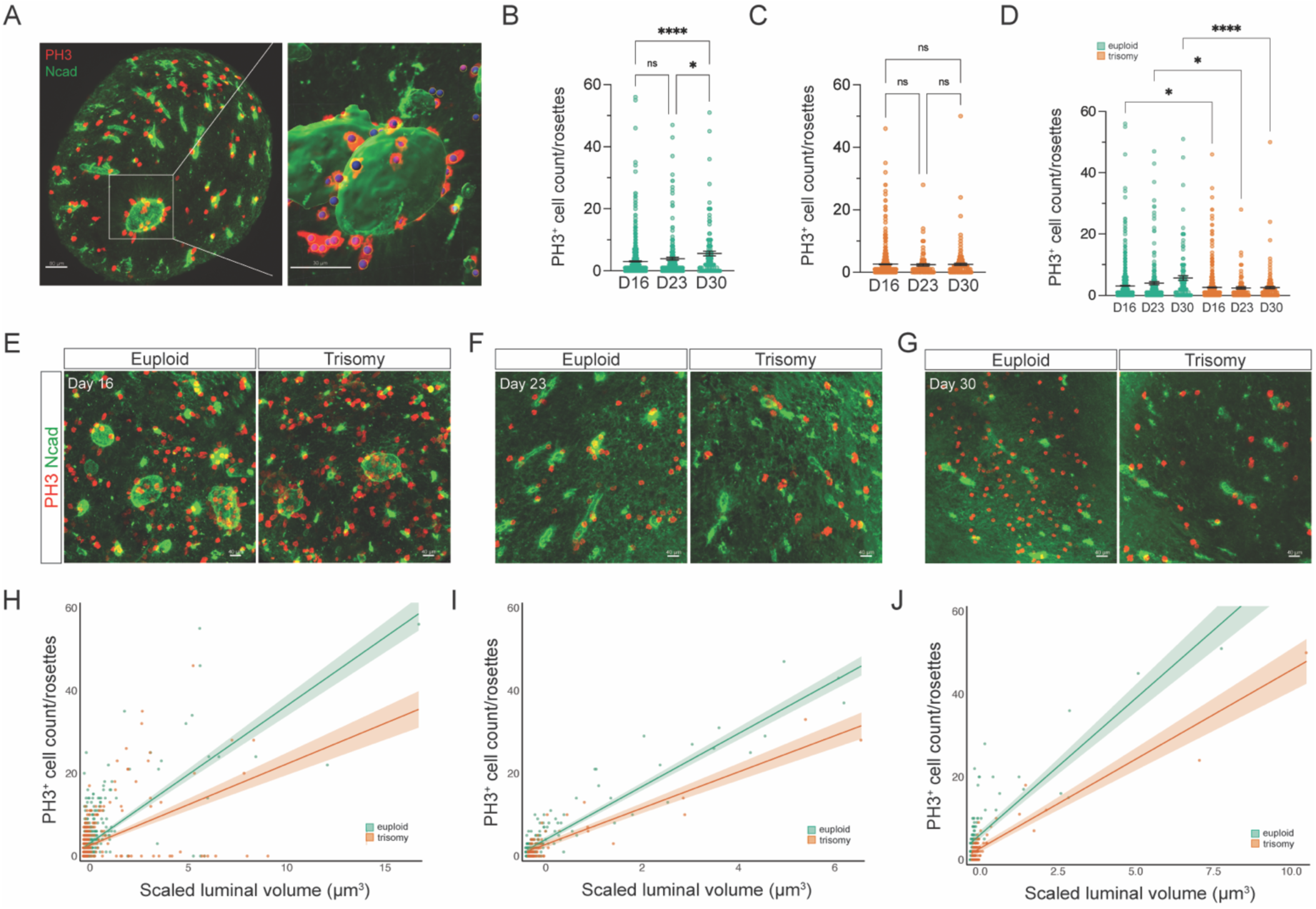
Mitotic activity in DFS rosettes. (**A**) Representative image of an Intact spheroid immunostained for PH3 and Ncad. Inset, higher- magnification view showing assignment of PH3^+^ spots to Ncad^+^ luminal surface using the spot-surface correlation function in imaris. **(B, C)** Quantification of PH3^+^ cells per rosette in euploid **(B)** and trisomic **(C)** spheroids at 16, 23, and 30 DIV. Data are shown as mean ± standard error of the mean (SEM). Euploid spheroids: one-way ANOVA test F = 9.383, *p* < 0.0001. **(D)** Comparison of PH3^+^ cells per rosette between genotypes at each timepoint. Data are shown as mean ± SEM; two-way ANOVA test F (5, 1382) = 8.063, *p* < 0.0001. **(E-G)** Representative images of euploid and trisomic rosettes at 16 **I**, 23 **(F)** and 30 DIV **(G)**. **(H-J)** Relationship between scaled luminal volume and PH3^+^ cell count per rosette at 16 **(I)**, 23 **(J)**, and 30 DIV **(K)**, showing that this relationship was attenuated in trisomic spheroids. Data collected from three differentiations, with 9 spheroids per genotype per timepoint. Statistical analyses performed using linear mixed-effects modeling followed by Type III analysis of covariance (ANCOVA). Data points represent individual rosettes. Statistical significance indicated as \**p* ≤ 0.05 and \*\*\*\**p* ≤ 0.0001. Scale bars, as indicated. PH3, red; Ncad, green.

Cumulative thymidine analog labeling has been used extensively *in vivo* to measure cell cycle dynamics within neural progenitor cell populations (Nowakowski et al., 1989; Takahashi et al., 1995). Because NPCs within rosettes are organized in a manner similar to the developing neural tube, this approach was adapted to quantify human NPC cycle dynamics within intact DFSs. Cohorts of nine spheroids per genotype and timepoint, across three independent differentiations were exposed to EdU for 24 hours to label progenitor cells during S-phase of the cell cycle. Subsets of each cohort were collected after 2, 4, 8, and 24 hours of EdU exposure and processed as whole mounts (**Figure 6A**). **Figure 6B** illustrates the expected cell-cycle distribution of EdU-labeled cells during cumulative labeling. After 4 hours of EdU exposure, NPCs in S-phase are labeled with EdU, while cells in M-phase are identified by PH3. By 8 hours, a subset of EdU-labeled progenitors has progressed into mitosis, resulting in colocalization of EdU and PH3 in mitotic cells. By 24 hours, all proliferating cells within the progenitor zone are expected to be EdU^+^ as unlabeled NPCs progress through S-phase during this time.

**Figure 6.**
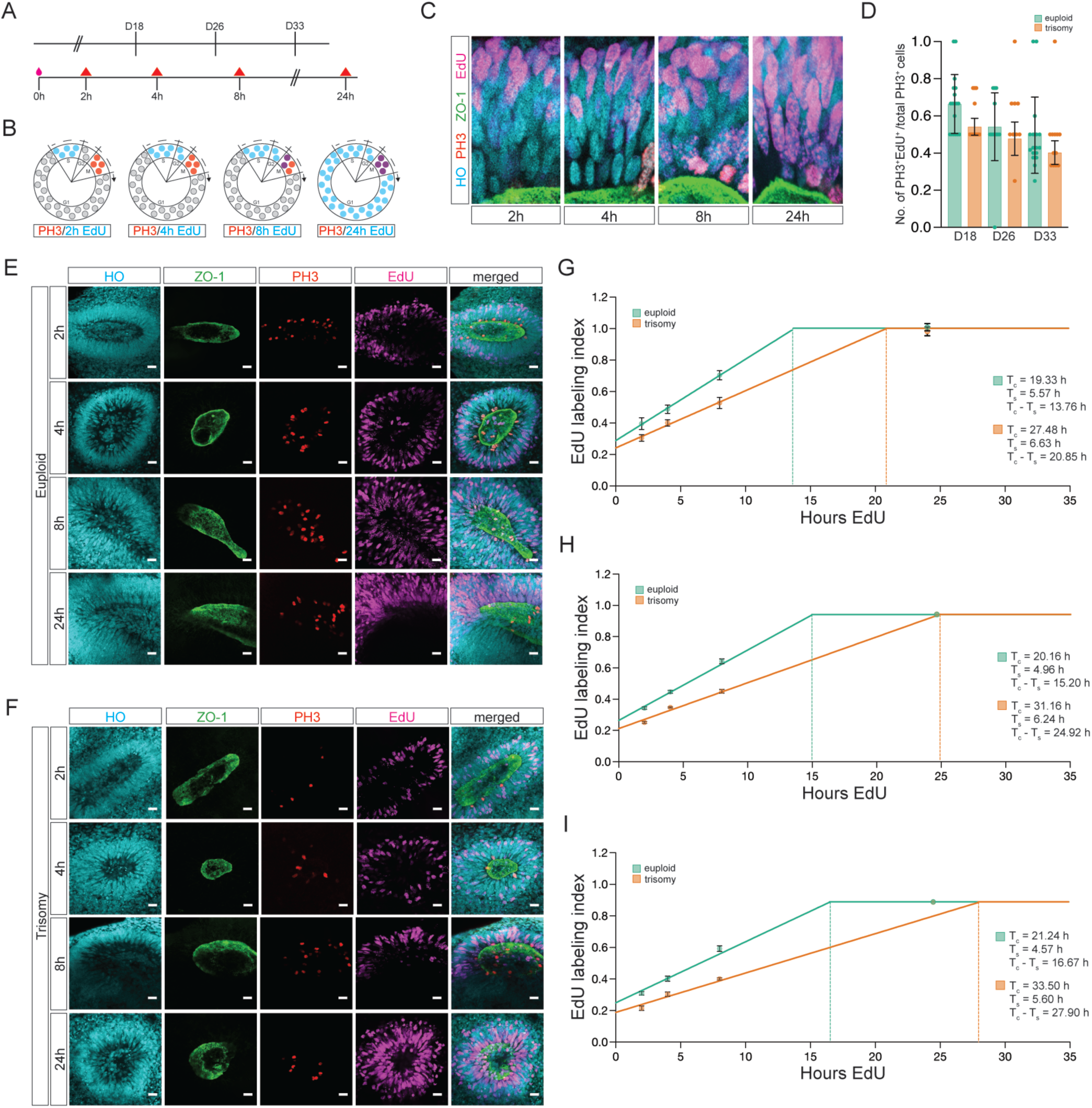
EdU labeling as a measure of cell cycle progression in DFS NPCs. **(A)** Schematic of cumulative EdU-labeling paradigm. Cohorts of spheroids were exposed to EdU for 24 h, and subsets were collected after 2, 4, 8, and 24 h of labeling. **(B)** Schematic depicting the expected cell-cycle distribution of EdU-labeled progenitors after 2, 4, 8, and 24 h of cumulative labeling. **(C)** High-magnification images illustrating interkinetic nuclear migration overt time. **(D)** Quantification of the proportion of PH3^+^/EdU^+^ cells after 8 h of EdU labeling in euploid and trisomic spheroids across developmental timepoints. **(E, F)** Representative images of rosettes following cumulative EdU labeling at 26 DIV, shown as single-channel and merged images for HO, ZO-1, PH3 and EdU. Images illustrate the progressive increase in EdU^+^ cells over the labeling interval. **(G-I)** Cumulative EdU-labeling curves used to estimate total cell cycle length (*T*_c_), S-phase duration (*T*_s_), and the non-S-phase components of the cell cycle (*T*_c_ – *T*_s_) in euploid and trisomic rosettes at 18 **(G)**, 26 **(H)**, and 33 DIV **(I)**. The slope of the line and the time required to reach the asymptote (growth fraction) were used to derive cell cycle parameters (Nowakowski et al., 1989; Takahashi et al., 1995). Data are shown as mean ± SEM. Scale bars, as indicated. HO, blue; ZO-1, green; PH3, red; EdU, magenta.

Our results demonstrate that several features of the *in vivo* ventricular zone organization are recapitulated in the rosette NPCs, including interkinetic nuclear migration (INM), characterized by spatially ordered progression of the nucleus position through the cell cycle phases. NPCs at the basal surface of the rosette incorporate EdU during S-phase and then their nuclei migrate toward the apical surface, where they undergo mitosis during M-phase (**Figure 6C**). Thus, rosettes recapitulate key spatial and proliferative properties of NPC dynamics *in vivo*.

We first quantified the fraction of PH3^+^ mitotic progenitor cells that were EdU^+^ after 4, 8, and 24 hours of EdU exposure. The greatest proportion of PH3^+^/EdU^+^ cells was found after 8 h of EdU exposure, suggesting that most EdU+ cells enter M phase within 8 hours after incorporating EdU. At each spheroid age examined, the proportion of PH3^+^/EdU^+^ cells at 8 h was lower in trisomic spheroids than in euploid controls (**Figure 6D**). Specifically, we find that 68.1% of euploid and 56% of trisomic NPCs were PH3^+^/EdU^+^ at 8 hours, suggesting changes in cell cycle kinetics as early as 18 DIV. Since these data indicate differences in the rate of NPC progression between phases of the cell cycle (S-phase to G2-phase), we turned next to a wholistic cell cycle measurement.

To estimate overall cell cycle phase durations and division rates of NPC within organized rosettes, the EdU labeling index was calculated as the percentage of EdU^+^ cells relative to the total number of cells within a defined rosette region of interest. From the series of images in **Figures 6E-F**, we observed that the EdU labeling index increased with exposure time, indicating continuous entry of unlabeled NPCs into S-phase. Linear regression of the cumulative labeling curves was then used to estimate total cell cycle length (T*_c_*) and S-phase duration (T*_s_*) (**Figures 6G-I**; **Table 3**). EdU experiments conducted on euploid spheroids at different DIV show that NPC cycle length progressively increased from 19.3 to 21.2 hours between 18–30 DIV, consistent with the average lengthening of the cell cycle during mammalian cortical development (Betizeau et al., 2013; Kornack and Rakic, 1998; Lukaszewicz et al., 2005; Takahashi et al., 1995).

**Table 3.** Summary of estimated cell cycle parameters in euploid and trisomic NPCs. Cell cycle phases were estimated from cumulative EdU-labeling analyses of rosette NPCs at 18, 26, and 33 DIV. Values reported in hours (h). Estimates generated from *n* = 6 rosettes from 2 spheroids per genotype at each developmental timepoint

| **DIV** | **Genotype** | **GF** | ***T*<sub>C</sub> – *T*<sub>S</sub> (h)** | ***T*<sub>S</sub> (h)** | ***T*<sub>C</sub> (h)** |
|---|---|---|---|---|---|
| D18 | Euploid | 1 | 13.76 | 5.57 | 19.33 |
|  | Trisomy | 1 | 20.85 | 6.63 | 27.48 |
| D26 | Euploid | 0.94 | 15.20 | 4.96 | 20.16 |
|  | Trisomy | 0.94 | 24.92 | 6.24 | 31.16 |
| D33 | Euploid | 0.88 | 16.67 | 4.57 | 21.24 |
|  | Trisomy | 0.88 | 27.90 | 5.60 | 33.50 |

*GF, Growth fraction; T<sub>S</sub>, duration of S-phase; T<sub>C</sub>, duration of cell cycle.*

The same cumulative labeling analysis was also applied to trisomic DFSs. Compared to their euploid counterparts, trisomic NPCs exhibited prolongation of the total cell cycle accompanied by increases in the combined G2+M+G1 (T*_c_* - T*_s_*) durations (**Figures 6G-I**). Specifically, the total NPC cycle increased from 27.5 to 33.5 hours, while the combined G2+M+G1 phase durations increased from 20.9 to 27.9 hours between 18–30 DIV. These findings indicate that trisomic NPCs progress more slowly through the cell cycle than euploid NPCs and that delayed proliferative kinetics is not restricted to a single phase of the cell cycle. Together with the slower growth of the PH3^+^ mitotic cell population, the EdU study uncovered significantly altered proliferative kinetics in trisomic NPCs from the expected euploid developmental trajectory.

To our knowledge, this approach represents one of the first applications of cumulative labeling analysis to quantify human NPC cell cycle kinetics within intact, spatially organized iPSC-derived neural rosettes. In addition, cumulative EdU-labeling assays also provided a more comprehensive measure of NPC proliferative behavior than PH3 or Ki67 imaging alone, allowing us to assess changes in NPC cycle kinetics in addition to transient measures of proliferation. This detailed analysis therefore elucidates potential cell cycle regulatory proteins and pathways that may be altered in trisomic NPCs. Together, these findings demonstrate the sensitivity and consistency of our methodology to uncover features of early human brain development and distinguish altered developmental trajectories associated with Down syndrome.

## DISCUSSION

This work establishes a rigorous experimental and methodological framework for future studies using programmed cortical spheroids to model quantitative features of typical brain development and the consequences of neurodevelopmental disorders. The methods allow standardized examination of complex developmental processes present at gross, structural, and cellular levels by implementing longitudinal experimental design with high-resolution 3D imaging and high-throughput image analysis.

A major strength of this study is the incorporation of analytical approaches that directly address known sources of variability in 3D culture systems. To address this variance, we used large sample sizes, longitudinal analysis, and distribution-based approaches, which enabled robust analysis of the time-dependent changes in developmental processes. Here we also used an isogenic trisomy 21 line to comprehensively test whether our approaches overcome the variability in the two compared experimental groups. Analysis of independent differentiations with multiple technical replicates demonstrated that the structural and cellular phenotypes were reproducible and enabled the identification of features that have not been reported in previous studies of Down syndrome or typical brain development. Specifically, we identified multimodal spheroid growth behavior and showed how a smaller spheroid subpopulation contributed to the overall reduction in trisomic spheroid size, thereby uncovering an unexpected variation in the trisomy 21 spheroid population. Accordingly, the method can identify phenotypes that may be missed when analysis relies only on average measurements at a single endpoint (Czerminski et al., 2022; Sandoval et al., 2024).

Another major strength of the approach is preservation of spatial organization. Unlike previous studies of cryosectioned organoids, rosettes in this work are characterized in intact preparations using whole-mount optical clearing, preserving their architecture and spatial organization. Analysis of the intact spheroids enabled a more complete assessment of rosette abundance and morphology, including the internal structures that may be missed or separated from their spatial context in selected histological sections. Our methods implemented high-throughput quantification approaches that captured details of a substantial number of rosettes compared to previous organoid models. Therefore, whole-mount 3D imaging provides a comprehensive and spatially resolved assessment of rosette morphometry throughout the intact spheroid.

The isogenic pair of iPSC lines used in our experiments provides an important proof-of-principle for this framework. Because the trisomic and euploid lines are genetically identical apart from the presence of a third copy of human chromosome 21 (HSA21), observed phenotypic differences can be attributed to trisomy 21 rather than confounding genetic background variation (Maclean et al., 2012; Hibaoui et al., 2014; Murray et al., 2015; Weick et al., 2013). Using this system, we detected altered spheroid growth, disrupted rosette morphogenesis, impaired mitotic activity and prolonged NPC cycle progression in the trisomic spheroids. These data demonstrate that the method can characterize multiple interacting features of early neurodevelopment in both typical and altered neurodevelopmental conditions.

### Implications for Down syndrome brain development

While there is a general consensus between human and mouse model studies on the presence of early brain development alterations in Down syndrome, the precise nature of the underlying mechanisms leading to microcephaly and/or intellectual disability in people with DS have not been consolidated between models and human subjects. For example, while cell cycle lengthening has been shown in several trisomic mouse models (Chakrabarti et al., 2007; Contestabile et al., 2007, 2009), human studies have been relegated to staining for Ki67 and other mitotic markers. Here, for the first time, our 3D analysis and imaging methods have discovered a significant lengthening of the cell cycle in trisomic dorsal forebrain NPCs. Furthermore, we show that the time-dependent lengthening of the cell cycle duration occurs both in control and trisomic spheroids as it does in the brain, supporting the translational validity of the spheroid platform. This confirmation is a critical step in supporting future studies of brain development in Down syndrome and in confirming key aspects of both the mouse models and the 3D culture system.

In this study, we focused our examination on repetitive differentiation experiments using cells from a single donor. Future studies should determine whether cells from other individuals with Down syndrome exhibit similar quantitative differences in early spheroid growth and development. This will likely be best accomplished by comparison to other isogenic lines. This will help to determine whether the specific timing of spheroid growth divergence observed here represents a general neurodevelopmental feature of Down syndrome or may instead reflect donor-specific variation. Thus, future comparative work between individuals using the methods described herein will begin to address the underlying framework leading to variations in nervous system function amongst individuals with Down syndrome. Regardless, our demonstration that structural and cellular phenotypes are reproducible across independent differentiations supports the robustness of the framework and that the methodology can be used to test reproducibility across additional cell lines.

The significance of this methodology approach extends beyond Down syndrome. Many neurodevelopmental conditions are expected to alter early progenitor behavior, tissue organization, cell cycle dynamics or developmental timing (Calegari and Huttner, 2003; Dehay and Kennedy, 2007; Gough et al., 2020). However, these phenotypes may be subtle, heterogeneous, or sensitive to line-to-line and batch-to-batch variation (Czerminski et al., 2022; Sandoval et al., 2024). The combination of isogenic design, longitudinal analysis, intact 3D imaging, rosette scoring and cell-cycle quantification provides a strategy for detecting such phenotypes in a reproducible and biologically meaningful manner. Therefore, this framework can be applied to neurodevelopmentally altered cell lines to determine whether altered growth, rosette morphogenesis and neural progenitor cell dynamics represent shared or condition-specific mechanisms of altered early brain development.

## LIMITATIONS AND FUTURE DIRECTIONS

A central consideration in interpreting these findings is the extent to which spheroids faithfully model *in vivo* human cortical development. Despite their translatability, iPSC-derived spheroids lack an intact tissue environment and do not provide a behavioral readout, which limits their ability to replicate the full complexity of the developing neocortex, including vascularization, neural connectivity and organized lamination (Di Lullo and Kriegstein, 2017; Qian et al., 2019). However, cortical spheroids recapitulate key features of early neural progenitor cell dynamics, including neural tube-like cytoarchitecture, apical mitosis and interkinetic nuclear migration (Broccoli et al., 2014; Kadoshima et al., 2013; Lancaster et al., 2013). Therefore, the model is highly valuable for investigating early cellular and molecular consequences of altered neurodevelopment.

A major future direction is to integrate this structural and cellular workflow with molecular profiling. Bulk RNA sequencing would help determine when transcriptional differences emerge in altered spheroids and whether they coincide with or precede structural and cellular deficits. This is especially relevant as previous sequencing studies have identified developmentally dynamic transcriptional changes in Down syndrome that were likely to affect NPC proliferation, cell-cycle regulation, and neuronal differentiation (Letourneau et al., 2014; Olmos-Serrano et al., 2016; Sobol et al., 2019). Single-cell multiome sequencing could further identify perturbed progenitor subpopulations and changes in chromatin accessibility during early spheroid development (Vuong et al., 2026). In this way, the framework described here provides a foundation for linking molecular disruptions to causal mechanisms of brain function in a fully translatable model system.

## EXPERIMENTAL PROCEDURES

### Resource availability

Corresponding author

Further information and requests for resources and reagents should be directed to the lead contact, Tarik F. Haydar at.

### Materials availability

This study did not generate new unique reagents. The WC-24-02-DS isogenic iPSC pair was generated and validated by the Bhattacharyya lab at the University of Wisconsin Madison and banked at WiCell.

### Data and code availability

Image analysis files, segmentation parameters and R scripts used for statistical analysis will be made available upon request.

### Experimental model and subject details

Experiments were performed using an isogenic iPSC paired line derived from a 25-year-old woman with mosaic Down syndrome. The isogenic pair consists of a trisomic line WC-24-02-DS-M and euploid line WC-24-02-DS-B, which are genetically identical apart from the presence of a third copy of HSA21. iPSCs were maintained in mTeSR™ Plus media (StemCell Technologies, 100-0276) supplemented with Primocin® (InvivoGen, ant-pm-05) on Cultrex® (R&D Systems, 3434-010-02) coated plates. iPSCs used for each experiment were at passages 14-17 for the trisomic line and 20-23 for the euploid line. Cultures were routinely screened for mycoplasma contamination and periodically submitted for karyotype analysis to confirm the continued presence of trisomy 21 and the absence of other chromosomal abnormalities.

### Generation of dorsal forebrain spheroids from iPSCs

The WC-24-02-DS isogenic pair was differentiated three independent times to human cortical spheroids with dorsal forebrain characteristics in accordance with established protocols (Birey et al., 2017; Yoon et al., 2019), adapted using the STEMdiff Dorsal Forebrain Organoid Differentiation Kit (StemCell Technologies, 08620). Briefly, plated iPSC colonies were dissociated into single cells with EDTA (ThermoFisher Scientific, 15575020) and resuspended in Seeding medium. The single-cell suspension was then transferred to one well of an AggreWell™800 24-well plate (StemCell Technologies, 34850) at a density of 10,000 cells per microwell. From days 1-5, the medium was changed to Forebrain Organoid Formation medium, with partial media changes performed daily. On day 6, aggregates were harvested and transferred to 6-well Ultra-low Attachment plates in Forebrain Organoid Expansion medium, with no more than 40 aggregates seeded per well. From days 8-24, aggregates were maintained in Forebrain Organoid Expansion medium with full-medium changes every 2 days. From day 25 onward, the medium was changed to Forebrain Organoid Differentiation medium, with full-medium changes every 2 days. Aggregates or spheroids were maintained until 35 DIV.

### Spheroid size by phase-contrast microscopy

To assess DFS growth, phase-contrast images of live whole spheroids were captured every two days from 8-30 DIV using a Keyence BZ-X710 inverted fluorescence microscope equipped with a Plan Fluor 4×/0.20 NA objective. Spheroid area (mm^2^) was measured in QuPath image analysis software using pixel classification-based segmentation. Pixel size was calibrated prior to analysis to enable conversion of pixel area to mm^2^.

### Whole-mount immunocytochemistry and clearing

Spheroids were fixed and stained at 16, 23 and 30 DIV following protocols adapted from Dekkers et al. (2019) for whole-mount organoid immunostaining. Spheroids were transferred to a 24-well Ultra-low Attachment plate (Corning® Costar®, CLS3473) using a wide-bore 1000-μL pipette tip (USA Scientific, 1011-9410) coated with 1% (wt/vol) bovine serum albumin fraction V (BSA) (Gibco, 15260037) in 1× phosphate-buffered saline (PBS) (Gibco, 14190250) and fixed in 4% (wt/vol) paraformaldehyde (PFA) (Thermo Scientific, AAJ19943K2) in 1× PBS for 20–45 minutes, depending on spheroid size, at 4°C. Fixed spheroids were permeabilized, blocked and incubated with primary antibodies (2× concentration) in organoid washing buffer (OWB) blocking solution consisting of 0.2% (vol/vol) Triton® X-100 and 0.2% (wt/vol) BSA in 1× PBS at 4°C for 2-3 days with mild rocking. Spheroids were washed three times in OWB for 2 hours per wash, incubated with secondary antibodies (2× concentration) in OWB for 2-3 days. Following additional 2-hour washes, spheroids were then incubated with Hoechst (Invitrogen™, H21486, 1:1000) in 1× PBS for 5 minutes at room temperature. Spheroids were then cleared in fructose-glycerol clearing solution consisting of 2.5 M fructose (Sigma-Aldrich, F0127) and 60% (vol/vol) glycerol (Sigma-Aldrich, G5516) in distilled water at −20°C until further slide mounting. Incubation time required for clearing depended on the age of the spheroid, with day 8 spheroids requiring 12-24 hours and day 30 spheroids requiring 3 days or more of incubation. Cleared samples could be stored at −20°C for up to one month before slide mounting. Whole-stained spheroids were mounted on custom slides with low-melting point agarose to maintain spheroid position for imaging. Whole-stained spheroids were mounted on custom slides with 4% (wt/vol) UltraPure™ low melting point agarose (Invitrogen, 16520050) in 1× PBS to maintain spheroid position for imaging. Mounting required a specialized slide preparation method. Two rectangular coverslips (Fisherbrand™, 12-541-035) were fixed to both sides of an 8-9 mm diameter × 0.8 mm depth Silicone Isolator™ (Grace Bio-labs, 666505) to maintain spheroid morphology. Once mounted and cleared, slides were stored at −20°C for up to six months until imaging. Primary antibodies used were rabbit anti-Sox2 (1:500, Abcam, ab97959), mouse anti-N-cadherin (Ncad) (1:250, BD Biosciences, 610920), and rabbit anti-phospho-histone H3 (PH3) (1:250, Sigma-Aldrich, 06-570). Alexa Fluor secondary antibodies (Invitrogen, 1:250) used were donkey anti-rabbit 546 (A10040) and anti-mouse 488 (A21202).

### High-resolution imaging and 3D reconstruction

Whole-mount cleared spheroids were imaged using an Olympus FVMPE-RS multiphoton microscope operated through FV31S-SW software, equipped with 25×/1.05 NA water-immersion objectives. Excitation was achieved using a 920 nm IR laser (MaiTai e HP DS-OL) for the green channel and a 1120 nm IR laser (INSIGHT X3 DUAL-OL) for the red channel. For larger spheroids, tile scans were acquired and stitched using FV31S-SW software. For Z-stack acquisition, images at 1024×1024 resolution were taken at step intervals of 1 μm with the 25× objective. Imaris software (version 11.0.0) was used for 3D reconstruction and surface-based segmentation of Sox2^+^ and Ncad^+^ structures in multiphoton data. Briefly, surfaces were initially generated across the entire spheroid using machine learning-based segmentation to identify rosette structures. For deeper rosette, segmentation was performed within user-defined ROIs to improve detection accuracy. Segmentation was performed with a surface smoothing parameter of 2 μm. Generated surfaces were filtered by volume or area, followed by machine learning-based classification to refine object selection. Volume and surface area were quantified for all segmented surfaces for subsequent analysis.

### Cumulative EdU labeling and cell counting

To characterize cell cycle phase duration and division rate of NPCs within rosettes, 5-ethynyl 2′- deoxyuridine (EdU) proliferative assays were performed using Click-iT™ EdU Alexa Fluor 647 imaging kit (Invitrogen, C10340), according to the manufacturing instructions. EdU was added to Forebrain Organoid media at 18, 26 and 33 DIV to label dividing progenitor cells within spheroids. The EdU-containing medium remained on the spheroids for a total labeling period of 24 hours. Spheroids were collected after 2, 4, 8 and 24 hours in culture and processed as whole mounts as described in the immunocytochemistry section. EdU-labeled spheroids were co-stained with primary antibodies, mouse anti-ZO-1 (1:250, BD Bioscience, 610967) and rabbit anti-PH3 (1:250, Sigma-Aldrich, 06-570). Alexa Fluor secondary antibodies (Invitrogen, 1:250) were donkey anti-rabbit 546 (A10040) and anti-mouse 488 (A21202). Z-stacks and tile scans were acquired to capture up to six rosettes per whole-mount spheroid using an Olympus FV4000 confocal laser scanning microscope with a 30× silicone immersion objective, operated through Fluoview™ software. Labeled cells were manually counted using ImageJ/Fiji. Cell cycle parameters, including growth fraction (GF), total cell cycle length (Tc), duration of S-phase (Ts), and time to reach GF (Tc – Ts), were calculated using linear regression analysis (Nowakowski et al., 1989; Takahashi et al., 1995).

### Quantification and statistical analyses

All statistical analyses were performed in RStudio and GraphPad Prism. Given the hierarchical structure of the dataset, with multiple rosette-level measurements nested within spheroids across genotypes and differentiation timepoints, linear mixed-effects models were used to account for non-independence among observations. Residual diagnostics were used to assess normality and heteroscedasticity, which were addressed using data transformation and model specification where appropriate. Rosette parameters were log-transformed prior to analysis to account for non-normality and unequal variance. Post-hoc pairwise comparisons between genotypes were performed at each timepoint using estimated marginal means with Holm-adjusted p-values for multiple comparisons.

To assess the relationship between progenitor zone volume and luminal volume across genotypes, a mixed-effects ANCOVA model was fitted to the data. Progenitor zone volume was modeled as the response variable, luminal volume as a continuous covariate, genotype as a fixed categorical effect, and differentiation and spheroid identity included as random effects to account for variability among spheroids and differentiation batches. The same modeling framework was applied to evaluate the relationship between luminal volume and PH3+ cell counts per rosette. Model fit was assessed using residual-versus-fitted and Q-Q plots. Statistical significance of fixed effects was assessed using Type III ANOVA.

Additional analyses were performed in GraphPad Prism for confirmation and visualization purposes. A two-tailed unpaired Student’s t-test was used to compare rosette density between euploid and trisomic spheroids across timepoints. A two-way ANOVA with a post-hoc Sidak’s multiple comparisons test was used to compare changes in the number of PH3+ cells per rosette across genotypes and timepoints. One-way ANOVAs with Sidak’s post hoc tests were used to compare changes in PH3+ cells per rosette within each genotype separately. All significance values presented in the text are defined as p < 0.05.

## Acknowledgements

We are deeply grateful to the donor whose generous contribution of cells made this work possible and to Dr. Anita Bhattacharyya of the University of Wisconsin for generating the WC-24-02-DS-B/M isogenic iPSC lines. We extend our gratitude to Dr. Jyoti Jaiswal for providing access to confocal imaging instrumentation at the Children’s National Research & Innovation Campus. We acknowledge the members of the Haydar Lab for their support of this research.

## Author Contributions

A.A. and T.F.H. conceived the study. AA designed and performed all experiments, acquired data and conducted data analyses. L.L. contributed to data analysis and validation of machine-learning and manual surface-based segmentation in imaris. A.A. and T.H. wrote the manuscript. All authors reviewed the article and approved the submitted version.

